# Arabidopsis DXO1 affects the processing of precursors of cytoplasmic and chloroplast ribosomal RNA

**DOI:** 10.1101/2022.09.14.507922

**Authors:** Monika Zakrzewska-Placzek, Aleksandra Kwasnik, Michal Krzyszton, Anna Golisz-Mocydlarz, Joanna Kufel

**Affiliations:** Institute of Genetics and Biotechnology, Faculty of Biology, University of Warsaw, Pawinskiego 5a, 02-106 Warsaw, Poland; Laboratory of Seeds Molecular Biology, Institute of Biochemistry and Biophysics, Polish Academy of Sciences, Pawinskiego 5a, 02-106 Warsaw, Poland

**Keywords:** Arabidopsis thaliana, chloroplast, nucleolus, pre-rRNA processing, ribosome biogenesis, rRNA, stress response

## Abstract

Decapping 5’-3’ exoribonucleases from the DXO/Rai1 family, are highly conserved among eukaryotes and exhibit diverse enzymatic activities depending on the organism. The biochemical and structural properties of the plant DXO1 differ from the yeast and animal counterparts, which is reflected in the *in vivo* functions of this enzyme. Here we show that Arabidopsis DXO1 contributes to the efficient processing of rRNA precursors in both nucleolar/cytosol and chloroplast maturation pathways. However, processing defects in DXO1-deficient plants do not depend on the catalytic activity of the enzyme but rely on its plant-specific N-terminal extension. Our RNA sequencing analyses show that the *dxo1* mutation deregulates the expression of many ribosomal protein genes, most likely leading to inefficient or delayed pre-rRNA maturation. Strikingly, some of the observed molecular and morphological phenotypes of *dxo1* plants are suppressed by the knock-down of *XRN3*, providing evidence for functional interaction between DXO1 and XRN proteins.

**HIGHLIGHT:** Arabidopsis DXO1 protein regulates the expression of genes encoding ribosomal proteins and contributes to the correct processing of ribosomal RNA precursors.

## INTRODUCTION

Ribosomes are complex macromolecular factories, composed of ribosomal proteins (r-proteins, RPs) and four ribosomal RNAs (rRNAs) that catalyze the translation of mRNAs into proteins. The biogenesis of ribosomes is controlled at every stage, from rRNA transcription, through co-transcriptional pre-rRNA processing, to the assembly of rRNAs with ribosomal proteins. In eukaryotes, most protein synthesis occurs in the cytoplasm, however, mitochondria and chloroplasts contain their own ribosomes that produce a number of organellar proteins. Both cytoplasmic and organellar ribosomes are composed of the large (LSU) and small (SSU) subunits. The cytoplasmic ribosome is an 80S ribonucleoprotein complex composed of a large 60S subunit, that contains 5.8S, 25S and 5S rRNAs, and a small 40S subunit with 18S rRNA (Weis *et al*., 2015*a*; Tomecki *et al*., 2017; Sáez-Vásquez and Delseny, 2019). The ribosomal RNAs form the catalytic and structural core of the ribosome. Three rRNAs, 5.8S, 25 and 18S, are transcribed as a polycistronic precursor by RNA polymerase I (Pol I) in the nucleolus, whereas 5S rRNA is synthesized by Pol III from separate transcription units distributed throughout the nucleus (Weis *et al*., 2015*a*; Tomecki *et al*., 2017; Sáez-Vásquez and Delseny, 2019). The primary 35S pre-rRNA transcript contains external transcribed regions at both ends (5’- and 3’-ETSs) and spacers between mature rRNA sequences (ITSs 1 and 2), which are systematically eliminated by a concerted action of multiple endo- and exoribonucleases to release mature 5.8S, 25S and 18S rRNAs. Pre-rRNA maturation requires not only enzymatic activities but also the binding of many factors that affect the structure of rRNA precursors and ensure the correct spatial and temporal coordination of these events.

In plants, two alternative pre-rRNA processing pathways have been described (see Figure 1A). The major “ITS1-first” pathway is initiated by cleavage within the ITS1 and separates precursors destined for the large and small ribosomal subunits, while the minor “5’ETS-first” pathway starts with the removal of the 5’ETS (Weis *et al*., 2015*a*; Tomecki *et al*., 2017). Interestingly, in *Arabidopsis thaliana* mutants in the rRNA processing factor IRP7, a third plant-specific pathway has been described, which begins with cleavage within ITS2 (Palm *et al*., 2019; Sáez-Vásquez and Delseny, 2019). This suggests a high plasticity of rRNA maturation pathways in plants.

**Figure 1.**
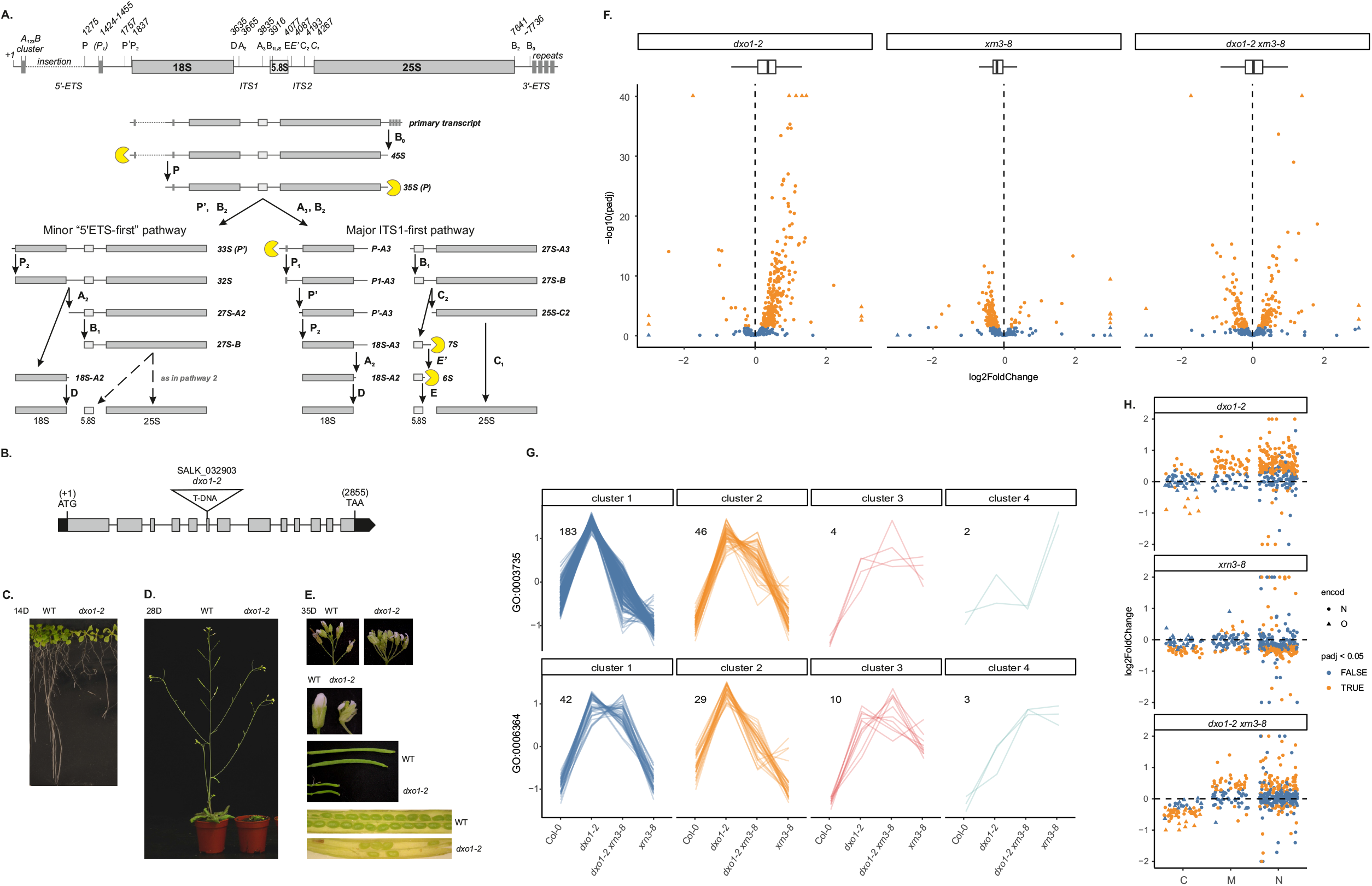
Phenotypes of *dxo1-2* plants are indicative of defects in ribosome biogenesis and chloroplast function. (A) Schematic representation of pre-rRNA structure and processing. (B) Structure of the *AtDXO1* (At4g17620) gene. Exons are represented by grey bars, the location of T-DNA insertion is indicated. (C-D) wild-type (WT) and *dxo1-2* plants grown on plates for 14 days (C) and in soil for 28 days (D). (E) Inflorescences, flower buds and siliques of 35-day-old plants. (F-H) Differential expression analysis by RNA-seq of ribosome-related genes between WT and mutant plants. Dots represent nuclear-encoded genes, squares represent mitochondrial-encoded genes and triangles represent genes encoded in organelles (F) Volcano plot and box plot of genes encoding structural constituents of the ribosome (GO:0003735). Orange shapes represent differentially expressed genes (padj < 0.05) and blue shapes represent non-differentially expressed genes, triangles mark out-of-scale genes. (G) Gene clustering analysis for genes with upregulated expression in the *dxo1-2* mutant and associated with Gene Ontology terms GO:0003735 (upper panel) or GO:0006364 (rRNA processing, lower panel). The number of genes in each cluster is shown on the diagram. Genes in each cluster are listed in Supplementary Table S1. Clustering was performed using standard R functions. (H) A scatter plot of genes encoding structural constituents of the ribosome (GO:0003735) divided into subgroups corresponding to the subcellular localization of the ribosome, C – chloroplast, M – mitochondrial and N – cytosolic ribosome. Orange represents differentially expressed genes (padj < 0.05) and blue non-differentially expressed genes. Dots and triangles correspond to nuclear-encoded (N) or organellar (O) genes, respectively.

To date, many ribosome biogenesis factors (RBFs) involved in rRNA maturation and modification in Arabidopsis have been identified. These include enzymes and ribonucleoprotein complexes that directly participate in pre-rRNA cleavage or trimming. The initial processing step in the 5’ETS at site P is carried out by the nucleolin-U3 snoRNP complex, while endoribonuclease NOB1 and the RNase III-type enzyme RTL2 are required for the proper P-A3 processing and cleavage in the 3’ETS, respectively (Comella *et al*., 2008; Kiyota *et al*., 2011; Missbach *et al*., 2013). The direct involvement of the remaining endonucleases characterized in yeast and mammals, i.e. RNase MRP, UTP24, LAS1 and RCL1, has not been demonstrated so far. In turn, exonucleolytic rRNA maturation by 5’-3’ exoribonucleases XRN2 and XRN3 includes the initial 5’ shortening of the primary transcript, trimming of the 27S-A3 and 26S processing intermediates and elimination of some excised pre-rRNA fragments (Zakrzewska-Placzek *et al*., 2010; Tomecki *et al*., 2017; Sáez-Vásquez and Delseny, 2019). Finally, the exosome exonuclease complex is responsible for the 3’ processing of several pre-rRNA species (Lange *et al*., 2011; Kumakura *et al*., 2013; Sikorski *et al*., 2015; Tomecki *et al*., 2017). Some of these enzymes most likely act redundantly; for example, knock-down of both XRN2 and XRN3 does not affect the level of mature rRNAs, suggesting the involvement of other exoribonucleases in the formation of rRNA 5’ ends (Zakrzewska-Placzek *et al*., 2010).

In chloroplasts, protein synthesis is carried out by the bacterial-type 70S chlororibosome composed of 50S LSU and 30S SSU. The large subunit contains three rRNA species, 23S rRNA, 5S rRNA and 4.5S rRNA (Yamaguchi and Subramanian, 2000; Ahmed *et al*., 2016; Bieri *et al*., 2017; Graf *et al*., 2017), whereas the small subunit is composed of 16S rRNA (Yamaguchi *et al*., 2000; Bieri *et al*., 2017; Graf *et al*., 2017; Perez Boerema *et al*., 2018). Although the composition and structure of chlororibosome have been studied using cryo-EM, less is known about the processing of chloroplast pre-rRNAs. Nonetheless, a number of factors implicated in rRNA maturation in chloroplasts have been identified in several plant species (Lu *et al*., 2011; Bang *et al*., 2012; Germain *et al*., 2013; Fristedt *et al*., 2014; Hotto *et al*., 2015; Liu *et al*., 2015; W *et al*., 2016). The chloroplast rRNA coding sequence is organized in an operon, transcription of which results in the synthesis of the polycistronic precursor containing all four chloroplast rRNAs and three tRNAs (16S-*trnI*-*trnA*-23S-4.5S-5S-*trnR*). Such pre-rRNA is subjected to endonucleolytic cleavages by RNase Z, RNase P and an unknown endonuclease, which together release tRNAs and separate rRNA precursors, and is followed by the formation of mature rRNA ends by exo- and endoribonucleases. These enzymes include exoribonucleases, RNase II (RNR1), RNase Mini-III and PNPase, and endoribonucleases, YbeY and RNase E, as well as RNase J, which has both exo- and endoribonucleolytic activities (Germain *et al*., 2013; Hotto *et al*., 2015; Liu *et al*., 2015).

The enzymes involved in pre-rRNA processing also include proteins from the DXO/Rai1 family, a small group of decapping enzymes implicated in essential mRNA surveillance mechanisms, namely mRNA 5’ end capping quality control (5’ QC) (Jiao *et al*., 2010, 2013; Chang *et al*., 2012) and removal of the non-canonical NAD^+^ cap (deNADding) (Fang et al., 2005; Gasse et al., 2015; Jiao et al., 2017; Kwasnik et al., 2019; Pan et al., 2020; Xue et al., 2000). Depending on the organism, DXO/Rai1 proteins have different properties and activities. *Schizosaccharomyces pombe* and *Saccharomyces cerevisiae* Rai1 exhibit two enzymatic activities: 5’ phosphohydrolase (PPH), which cleaves pyrophosphate from the 5’ triphosphate RNA, and non-canonical decapping towards unmethylated caps, removing the entire GpppN dinucleotide (Xiang *et al*., 2009; Jiao *et al*., 2010). Rai1 forms a heterodimer with a nuclear 5’-3’ exoribonuclease Rat1, which enhances/stabilizes Rat1 activity in several processes, including transcription termination and rRNA maturation (Xue *et al*., 2000; Xiang *et al*., 2009). Rai1-Rat1 together with the Las1-Grc3 heterodimer form a complex that functions in the 5’ trimming of the 26S intermediate released by Las1 cleavage at site C2 within ITS2 (Gasse *et al*., 2015; Fromm *et al*., 2017). Since the 26S precursor generated by Las1 has a 5’ hydroxyl group (5’-OH), its further exonucleolytic maturation requires phosphorylation by Grc3 kinase. Remarkably, yeast Rai1 has been shown to possess a new 5’ hydroxyl dinucleotide hydrolase (HDH) activity, potentially allowing the Rat1-Rai1 complex to efficiently degrade 5’-OH RNAs (Doamekpor *et al*., 2020*a*). LAS1-dependent processing steps have not been described in Arabidopsis, except for the nucleolar localization of the LAS1-GRC3 dimer (Maekawa *et al*., 2018; Sáez-Vásquez and Delseny, 2019).

Several fungal species have a second DXO/Rai1 family protein, Dxo1, which has no pyrophosphate activity but, in addition to removing unmethylated caps and deNADing, is able to completely degrade RNA owing to its 5’-3’ exoribonucleolytic activity (Chang *et al*., 2012; Jiao *et al*., 2017). Importantly, it has been shown recently that the primary role of Dxo1 in *S. cerevisiae* is the final 5’ end processing of 25S rRNA in the cytoplasm (Hurtig and van Hoof, 2022).

The mammalian DXO/Rai1 homolog, which is a functional hybrid of fungal Rai1 and Dxo1 enzymes, functions in 5’ QC, deNADding and removal of other non-canonical caps from RNA (Jiao *et al*., 2013; Doamekpor *et al*., 2020*b*), but its contribution to pre-rRNA processing has not been addressed. Unlike fungal Rai1, DXO most likely does not interact directly with the Rat1 homologs, the XRN proteins, as it lacks an appropriate binding domain. Arabidopsis DXO1, a single DXO/Rai1 family protein, shows remarkable differences in biochemical properties compared to its fungal and animal counterparts (Kwasnik *et al*., 2019; Doamekpor *et al*., 2020*a*). Although the active site of DXO proteins is highly conserved among eukaryotes, Arabidopsis DXO1 contains a single amino acid substitution near the active site that affects the properties of the enzyme. As a result, plant DXO1 has very limited 5’ QC and 5’-3’ exoribonuclease activities, but is a potent deNADding enzyme (Kwasnik *et al*., 2019; Pan *et al*., 2020). Importantly, although DXO1 is not present in chloroplasts, *dxo1* mutants exhibit phenotypes related to chloroplast function, including low chlorophyll content and decreased levels of chloroplast-encoded mRNAs (Kwasnik *et al*., 2019; Pan *et al*., 2020).

In this work, we report that DXO1 is required for accurate and efficient processing of pre-rRNAs in Arabidopsis. Strikingly, DXO1 contributes not only to the maturation of rRNAs in the nucleolus but also in chloroplasts. The *dxo1* mutants have a pointed-leaf phenotype indicative of ribosome-related dysfunction, combined with retarded growth and reduced fertility, and respond differently to various stress factors, such as salinity and sugar deficiency or excess. On the molecular level, *dxo1* mutants exhibit a strong accumulation of several nuclear/nucleolar and chloroplast pre-rRNAs. Additionally, according to our RNA-seq data (Kwasnik *et al*., 2019), the *dxo1* mutation alters the expression of many genes encoding ribosomal proteins and ribosome biogenesis factors. We postulate that this is the likely source of the observed pre-rRNA processing defects.

## MATERIALS AND METHODS

### Plant material and growth conditions

*Arabidopsis thaliana* wild-type, mutant and transgenic plants used in the study are in the Columbia-0 (Col-0) genetic background. We used the following mutant lines: *dxo1-2* (SALK_032903), *xrn2-3* (SALK_114258), *xrn3-8* (RNAi-silenced line (Zakrzewska-Placzek *et al*., 2010)) and transgenic *dxo1-2* lines expressing DXO1 variants: DXO1 (E394A/D396A), DXO1 (ΔN194), DXO1 (ΔN194 / E394A / D396A) (Kwasnik *et al*., 2019). The *xrn2-1 xrn3-3* mutant line (Gy *et al*., 2007) was obtained from Allison Mallory. The *dxo1-2 xrn2-3* and *dxo1-2 xrn3-8* double mutant lines were generated by crossing the corresponding single mutant lines, the *dxo1-2 xrn2-3 xrn3-8* triple mutant was constructed by crossing the *dxo1-2 xrn3-8* line with the *xrn2-3* mutant. Unless stated otherwise plants were grown in Percival growth chambers at 22/19°C on a 16-h-light and 8-h-dark (long-day) photoperiod. Seeds were germinated in soil or were surface-sterilized with 30% bleach/0.02% Triton-X100 solution and germinated on half-strength MS (Murashige and Skoog, 1962) basal salt medium supplemented with 1% (w/v) sucrose and 0.3% phytagel (Sigma-Aldrich).

### Stress resistance tests

For stress treatments seeds were surface sterilized and grown on MS (Murashige and Skoog, 1962) medium supplemented with 1% (w/v) sucrose and 0.3% phytagel under long-day conditions, then after 2 weeks seedlings were subjected to high temperature, drought or wounding. Samples were collected at the indicated time points, blotted dry and immediately frozen in liquid nitrogen. In the case of high-temperature stress, plates were transferred to a 37°C growth chamber and samples were collected after 10, 20 and 30 min. For drought stress, seedlings were picked from plates, weighed and dried at RT for 30 or 60 min. Wounding was performed by crushing with brass kenzan. Each plate was crushed 10 times and samples were taken after 15 and 30 minutes. For salt and glucose treatments, seeds were grown on MS (Murashige and Skoog, 1962) medium containing NaCl and 1% (w/v) sucrose or only glucose (without sucrose). In both cases the media contained 0.3% phytagel (Sigma-Aldrich). The appropriate amounts of sterilized NaCl or glucose solutions were added to cooled autoclaved media. The seeds were germinated on the NaCl- or glucose-containing plates for up to 21 days under long-day growth conditions. Samples collected from each stress treatment were subjected to RNA isolation and Northern blot analysis.

### RNA methods

<u>Total RNA extraction.</u> Total RNA was isolated from 14-day-old seedlings using Trizol reagent (Sigma-Aldrich) according to the manufacturer’s instructions. <u>Northern blotting.</u> Low-molecular-weight RNAs were separated on 6% acrylamide/7 M urea gels and transferred to a Hybond-XL membrane (GE Healthcare) by electrotransfer. High-molecular-weight RNAs were analyzed on 1.1% agarose/6% formaldehyde gels and transferred to a Hybond-XL membrane by capillary elution. γ-^32^P 5′ end-labelled oligonucleotides (Supplementary Table S2) were used as probes against precursor and mature RNAs and U2 snoRNA-specific probe as a loading control. Probes were end-labeled using T4 Polynucleotide Kinase (Thermo Scientific) with [γ-^32^P] (SRP-201, Hartmann Analytic). mRNAs were detected using random primed probes, amplified on a cDNA template with respective primers (Supplementary Table S2) and DECAprime™ II labeling kit (Thermo Scientific) and [α-^32^P] dATP (SRP-203, Hartmann Analytic). Quantification of northern blots was performed using Typhoon FLA 9000 Gel Imaging Scanner (GE Healthcare) and ImageQuant (GE Healthcare) or ImageJ (U. S. National Institutes of Health, https://imagej.nih.gov/ij/) software. <u>Primer extension</u> was performed as described (Zakrzewska-Placzek *et al*., 2010) using 10 µg of total RNA and γ-^32^P 5′-end-labelled primer 25S-5′. Sequences of oligonucleotides used in this study are listed in Supplementary Table S2.

### RNA-sequencing

RNA-seq analyses of the *dxo1-2* and *xrn3-8* mutants, including the procedure and bioinformatics analysis, were described in (Krzyszton *et al*., 2018; Kwasnik *et al*., 2019). The sequencing and bioinformatics analysis of the double mutant *dxo1-2 xrn3-8* was performed simultaneously using the same protocols. Libraries were paired-end sequenced on HiSeq4000 (DNA Research Centre, Poznan, Poland). RNA-seq data are deposited in the Gene Expression Omnibus database under accession codes GSE95473 (Col-0 and *xrn3-8* plants) and GSE99600 (*dxo1-2* plants). RNA-seq data for double *dxo1-2 xrn3-8* mutant and count data for all genotypes were deposited upon GSE210614. Library preparation protocol for 3’RNA-sequencing was described in (Krzyszton *et al*., 2022) and sequencing was performed at the core facility of the International Institute of Molecular and Cell Biology in Warsaw using the NextSeq 500. Data were analyzed as described earlier (Krzyszton *et al*., 2022). Briefly, reads R1 and R2 were processed separately. Read R1 fastq file was transformed using *awk ‘NR%2==1 {print $0} NR%2==0 {print substr($1,16,5) substr($1,1,15)}’*. Read R2 was trimmed to remove potential contamination with poly(A) tail using BRBseqTools (v 1.6) Trim (Alpern *et al*., 2019) and parameters *-polyA 10 -minLength 30*. Then mapped using STAR (v 2.7.8a) (Dobin *et al*., 2013) to TAIR 10 genome version and Araport11 genome annotation with parameters *--sjdbOverhang 54 --outSAMtype BAM SortedByCoordinate -- outFilterMultimapNmax 1*. Finally, the count matrix for each seed and each gene was obtained using BRBseqTools (v 1.6) CreateDGEMatrix with parameters *-p UB -UMI 14 -s yes*, using Araport11 genome annotation and a list of barcodes. Count matrixes from different libraries were combined and used for further analysis using DESeq2 (Love *et al*., 2014). Data for 3’RNA-seq are deposited upon GSE210631. Genes belonging to GO:0003735 and GO:0006364 GO terms were obtained from plant.enesembl.org. GO term enrichment analysis was done using the PANTHER 17.0 Overrepresentation Test (GO Ontology database DOI: 10.5281/zenodo.6799722 Released 2022-07-01) (Thomas *et al*., 2022) with Fisher’s extract test and Bonferroni correction for P<0.05.

## RESULTS

### Developmental defects of plants lacking DXO1 suggest its involvement in ribosome biogenesis and chloroplast functioning

To investigate the possible role of DXO1 in ribosome biogenesis we used the *dxo1-2* (SALK 032903) mutant line with a T-DNA insertion in the fifth intron of the AT4G17620 gene (Kwasnik *et al*., 2019) (Fig 1B). The *dxo1-2* plants have a chloroplast-associated phenotype with pale-green leaves and reduced chlorophyll content, most likely related to the downregulation of chloroplast-encoded mRNAs (Kwasnik *et al*., 2019). Importantly, the *dxo1-2* line also exhibits several phenotypic features that are similar to those found in ribosome-related Arabidopsis mutants. These include developmental defects and reduced fertility, as manifested by shortened siliques and a significant number of aborted seeds (Fig. 1C-E). The leaves of *dxo1-2* seedlings are narrow and pointed, and the primary roots are relatively short (Fig. 1C and Supplementary Fig. S1A), which resembles phenotypes observed in mutants of factors involved in ribosome biogenesis, namely *brx1* (Weis *et al*., 2015*b*), *rpl4* (Rosado *et al*., 2010), *atprmt3* (Hang *et al*., 2014), *apum23* (Abbasi *et al*., 2010; Huang *et al*., 2018), *mtr4* (Lange *et al*., 2011), *dim1A* (Wieckowski and Schiefelbein, 2012), *rid2* (Ohbayashi *et al*., 2011), *rh7* (Liu *et al*., 2016), some *oli* lines (Fujikura *et al*., 2009) and nucleolin mutants (Kojima *et al*., 2007; Petricka and Nelson, 2007). Phenotypes of *dxo1-2* plants and our previous data (Kwasnik *et al*., 2019), suggest that DXO1 is potentially involved in ribosome biogenesis and chloroplast function in Arabidopsis. The role of yeast DXO1 counterparts in pre-rRNA processing also supports this possibility (Fang et al., 2005; Gasse et al., 2015; Hurtig and van Hoof, 2022; Schillewaert et al., 2012; Xue et al., 2000) reviewed in (Tomecki *et al*., 2017)).

We previously demonstrated that the phenotype of the *dxo1-2* mutant was complemented by the expression of both the wild-type and the catalytically inactive DXO1 variants (E394A/D396A), suggesting that the enzymatic activity of DXO1 is either redundant with other enzymes or irrelevant to the phenotypes observed (Kwasnik *et al*., 2019). In contrast, *dxo1-2* lines expressing the N-terminally truncated forms, namely catalytically active DXO1 (ΔN194) and inactive DXO1 (ΔN194/E394A/D396A), retained the characteristic *dxo1-2* phenotype, indicating the important role of the DXO1 plant-specific extended N-terminus for its function in Arabidopsis (Kwasnik *et al*., 2019).

To characterize the role of DXO1 in ribosome biogenesis, alongside the *dxo1-2* mutant we also used *xrn2-3, xrn3-8* and double *xrn2-1 xrn3-3* mutants in the two Rat1 homologs, the nuclear 5’-3’ exoribonucleases XRN2 and XRN3, which show defects in pre-rRNA processing (Zakrzewska-Placzek *et al*., 2010). To investigate possible functional interconnections between DXO1 and nuclear XRN proteins, we generated double *dxo1-2 xrn2-3, dxo1-2 xrn3-8* and triple *dxo1-2 xrn2-3 xrn3-8* lines, which retained the morphological phenotype of *dxo1-2*, including pale green coloration, but showed a less severe growth defect than the single *dxo1-2* line (Supplementary Fig. S1A). Similarly, *dxo1-2 xrn3-8* and *dxo1-2 xrn2-3 xrn3-8* plants grew better than the single *xrn3-8* mutant. The phenotypes of double and triple mutants point to a partial genetic suppression of *dxo1-2* and *xrn3-8* by mutations in *XRN2/3* and *DXO1* genes, respectively, and provide evidence for functional interactions between DXO1 and XRN2/3 proteins.

### Mutation in *DXO1* alters the expression of many ribosome-related genes

To assess the transcriptional impact of DXO1 on mRNAs related to pre-rRNA processing and ribosome assembly, as well as the structure and function of chloroplasts, we used our earlier RNA sequencing (RNA-seq) data for *dxo1-2* (Kwasnik *et al*., 2019). This analysis revealed that a large number of genes was affected in the *dxo1-2* transcriptome and the set of upregulated genes showed a remarkable enrichment of GO-terms related to the ribosome and ribosome biogenesis, including ribosome, preribosome, ribosome assembly, rRNA processing, nucleolus, snoRNA binding, ribonucleoprotein complex, and translation pre-initation complex (Kwasnik *et al*., 2019). We have now performed a more detailed analysis of this particular set of misregulated genes associated with ribosome biosynthesis and function. Out of 4,878 genes with significantly increased expression, 4.8% (235 genes) corresponded to genes encoding ribosome structural components (GO:0003735, Fig. 1F), which is a relatively small number in the context of all altered genes. It is striking, however, that 55% of Arabidopsis genes belonging to this category have a higher expression level in the *dxo1-2* mutant compared to the wild-type. The majority of significant changes in ribosome-related mRNAs in *dxo1-2* do not exceed the 2-fold value, possibly because of their key cellular functions, limiting large alterations in the expression of these essential genes.

Considering the phenotypic suppression in plants lacking both DXO1 and XRN3, we compared the RNA-seq data of the *dxo1-2 xrn3-8* double mutant with the results of the corresponding single mutant lines and wild-type plants, obtained in the same RNA-seq experiment and already published (Krzyszton *et al*., 2018; Kwasnik *et al*., 2019). Gene clustering analysis revealed that a large number of genes affected in *dxo1-2* were also altered in *xrn3-8*. For many gene clusters, these genes had an intermediate level of expression in the double mutant, and in the case of some clusters similar to that in the wild-type. Genes with the restored or partially restored expression belong to a wide range of GO categories (PANTHER Overrepresentation Test (Thomas *et al*., 2022); Supplementary Table S1), including rRNA processing, ribosome maturation, assembly and function (Supplementary Fig. S1B, clusters 3 and 4), regulation of transcription and RNA processing (Supplementary Fig. S1B, cluster 5), tissue development and cell division, cell cycle (Supplementary Fig. S1B, clusters 7, 9 and 11), response to stress, defense response (Supplementary Fig. S1C, clusters 2, 10, 13 and 16), and finally signal transduction, transport and localization (Supplementary Fig. S1C, clusters 5 and 16).

Of particular interest are GO:0003735 (ribosome structural components) and GO:0006364 (rRNA processing) categories related to ribosome biogenesis and function (Fig. 1F-1G). For the GO:0003735 category we observed mostly opposite gene expression profiles in *dxo1-2* and *xrn3-8* mutants (Fig. 1F). Of the 235 genes annotated as structural constituents of the ribosome that were upregulated in *dxo1-2*, 183 had downregulated or unaffected expression in *xrn3-8* and averaged in the double mutant (Fig. 1G, upper panel and Supplementary Table S1). In turn, the GO:0006364 category revealed a slightly different pattern, in which expression of only 29 out of 84 genes upregulated in *dxo1-2* was at least partially attenuated by the additional *xrn3-8* mutation (Fig. 1G, lower panel, cluster 2 and Supplementary Table S1). Notably, expression profiles of genes affected in *dxo1-2* show that suppression in the double mutant does not apply to all clusters (Supplementary Fig. S1B-C and Supplementary Table S1), suggesting that this is not a general feature but occurs for specific types of genes. These observations are consistent with the partial suppression of *dxo1-2* mutation by *XRN3* silencing, but also suggest that different phenotypes may be affected to varying degrees.

Interestingly, the expression of only a small number of genes encoding structural components of the ribosome (GO:0003735) was decreased in the *dxo1-2* mutant. Of these, about 30% is encoded in the chloroplast genome (Fig. 1H), which is in agreement with the *dxo1-2* phenotypes associated with photosynthesis and chloroplasts and underlines the importance of DXO1 in regulating chloroplast activity ((Kwasnik *et al*., 2019) and this work). The impact of DXO1 on components of the cytoplasmic and chloroplast ribosome pathways raises the question of how DXO1 regulates ribosome biogenesis in these different cellular compartments.

### DXO1 is required for efficient processing within ITS2 of nuclear-encoded pre-RNA

Considering that many stages of ribosome biogenesis occur simultaneously and are interdependent (Peña *et al*., 2017), widespread changes in the expression of genes encoding ribosome components and biogenesis factors in the *dxo1-2* mutant may cause disruptions in rRNA maturation. The yeast Rai1 protein, as a part of the Las1-Grc3-Rat1-Rai1 complex, has been reported to be involved in the removal of the ITS2 spacer from the rRNA precursor (Gasse *et al*., 2015). To test whether Arabidopsis DXO1 also contributes to ITS2 processing, we performed a northern blot analysis of RNA extracted from *dxo1-2* plants, using a set of probes that recognize specific pre-rRNAs (Fig. 2 and Supplementary Fig. S2A). To assess the functional interconnections between DXO1 and nuclear XRN proteins at the molecular level we used *dxo1-2 xrn2-3, dxo1-2 xrn3-8* and *dxo1-2 xrn2-3 xrn3-8* mutants and, for comparison, *xrn2-1 xrn3-3* line, which presents a characteristic pattern of rRNA precursors (Zakrzewska-Placzek *et al*., 2010). Northern blot analysis revealed that ITS2 processing is indeed disturbed in the *dxo1-2* mutant. In particular, probes P5 and P44 located within ITS2 showed substantial accumulation of 27SB and 7S pre-rRNAs in *dxo1-2* plants, possibly reflecting ineffective ITS2 elimination in the absence of DXO1. This was accompanied by a modest upregulation of the 27S-A2 and 27S-A3 species, as shown by probes P42 and P4 (Fig. 2A). These effects, especially the increased level of 27SB and 7S precursors, were observed in *xrn2, xrn3* and all *dxo1* lines suggesting that pre-rRNA processing defects in plants lacking DXO1 and XRN proteins may have a common background.

**Figure 2.**
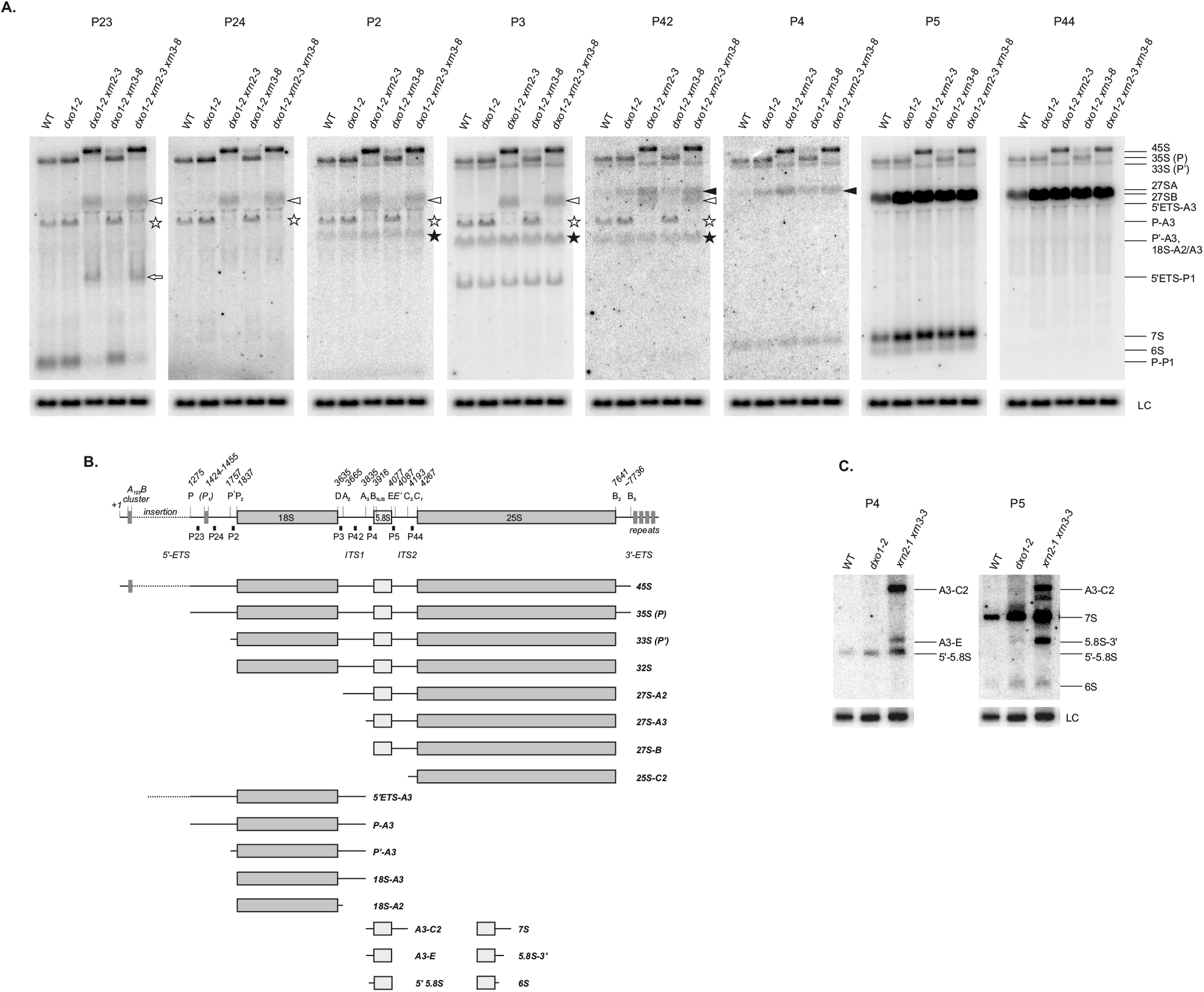
Lack of DXO1 results in the accumulation of several rRNA precursors. (A and C) Northern blot analysis of pre-rRNA precursors in WT, *dxo1-2, dxo1-2 xrn2-3, dxo1-2 xrn3-8* and *dxo1-2 xrn2-3 xrn3-8* mutants. Total RNA was extracted from 14-day-old seedlings and separated on 1.1% agarose (A) or 6% polyacrylamide (C) gels and hybridized with probes specific for particular regions of pre-rRNA depicted in (B). Hybridizations for U2 snoRNA were used as loading controls (LC). Identified pre-rRNAs are marked on the right. White arrowheads indicate the 5’ETS-A3 intermediate. Black arrowheads correspond to 27SA pre-rRNA. White asterisks indicate P-A3 and black asterisks denote P’-A3 and 18S-A2/A3 precursors. (B) Diagram showing the pre-rRNA processing intermediates and precursors detected by Northern blot analysis and showing the positions of hybridization probes. Sequences of hybridization probes are listed in Supplementary Table S2.

Although ITS2 maturation is not well characterized in Arabidopsis, LAS1 and GRC3 have been postulated to be involved in this process (Maekawa *et al*., 2018; Sáez-Vásquez and Delseny, 2019). Assuming that DXO1 plays an analogous role to Rai1 in ITS2 removal, the *dxo1-*2 mutation would lead to the accumulation of the 25S-C2 intermediate. However, northern hybridization with the P44 probe did not detect this species in *dxo1-2* (Fig. 2A and Supplementary Fig. S2A). Instead, we observed a faint band of a similar size, which may represent the 25S-C2 species, in the double *xrn2-1 xrn3-3* mutant (Supplementary Fig. S2A, probes p5 and p44). This suggests that in Arabidopsis XRN2 and XRN3, but not DXO1, may be involved in the elimination of ITS2 spacer and maturation of the 5’ end of 25S rRNA (Fig. 2A and Supplementary Fig. S2). The two XRN 5’-3’ exoribonucleases act redundantly in this process, as single *xrn2* or *xrn3* mutations are not sufficient to cause accumulation of 25S-C2 (Supplementary Fig. S2). Also checking for defects in 25S 5’ end maturation by primer extension did not show any differences in *dxo1-2* (Supplementary Fig. S2B). This is in contrast to the recently reported function of Dxo1 in yeast, where this enzyme performs the final trimming of 25S rRNA 5’ end (Hurtig and van Hoof, 2022).

Furthermore, hybridization with the P5 probe revealed that 5.8S pre-rRNAs are elevated in *dxo1-2* plants and double and triple mutants (Fig. 2A). To find out exactly what types of 5.8S precursors are affected in *dxo1-2* mutants, we performed northern blot analysis of WT, *dxo1-2* and *xrn2-1 xrn3-3* RNAs separated on polyacrylamide gels (Fig. 2C). As previously described (Zakrzewska-Placzek *et al*., 2010), 7S, 5’-extended 7S pre-rRNA (A3-C2), 6S, and 5’- and/or 3’-unprocessed 5.8S (A3-E, 5.8S-3’ and 5’-5.8S) species accumulated in the *xrn2-1 xrn3-3* mutant. In contrast, mainly 7S pre-rRNA and to a much lesser extent 6S, 5.8S-3’ and 5’-5.8S intermediates were increased in the *dxo1-2* mutant. Taken together, this analysis shows that *dxo1* mutation leads to the accumulation of normally occurring precursors rather than unusual or aberrant species, which is in agreement with the possible delay in ITS2 processing in the absence of DXO1. However, unlike the yeast Rai1/Rat1 complex, DXO1 is not involved in the formation of mature 5’ ends of 25S and 5.8S rRNAs but contributes, probably indirectly, to 5.8S rRNA 3’-end processing. The 3’-extended 5.8S precursors have been observed in mutants of yeast Rat1, Rrp17 or Rai1, in human cells deficient in XRN2 and Arabidopsis *xrn2* mutants (Fang et al., 2005; Schillewaert et al., 2012; Zakrzewska-Placzek et al., 2010). Our results support the hypothesis that the maturation of 5’ and 3’ ends of 5.8S rRNA by Rat1/XRN2 and the exosome is coordinated, which was proposed to result from the close physical proximity of 5.8S ends in the spatial structure of pre-60S particles (Schillewaert *et al*., 2012). Interestingly, ITS2 processing defects in the *dxo1-2* mutant were not attenuated by additional *xrn2* or *xrn3* mutations and vice versa (Fig. 2A, probes P5 and P44). This observation argues that mutual suppression of some molecular and morphological phenotypes does not apply to the maturation of nuclear pre-rRNAs. It is noteworthy that the expression of genes belonging to the GO:0006364 rRNA processing category that were altered in the *dxo1-2* mutant was restored to a very limited extent by *XRN3* silencing (see Fig. 1G, cluster 2).

Importantly, pre-rRNA processing defects observed in the *dxo1-2* mutant did not lead to changes in levels of mature ribosomal RNAs (Supplementary fig. S2C and D). A similar situation was reported for mutants in many plant rRNA processing factors (e. g.(Shi *et al*., 2005; Zakrzewska-Placzek *et al*., 2010; Abbasi *et al*., 2010; Lange *et al*., 2011)), probably due to the existence of alternative pathways and redundant enzymatic activities involved in plant rRNA biogenesis.

As in other eukaryotes, pre-rRNA processing machinery in plants also includes small nucleolar RNAs, some of which participate in pre-rRNA cleavages. The early cleavage at site P requires a U3 snoRNP complex containing U3 and U14 snoRNAs (Sáez-Vasquez *et al*., 2004; Pontvianne *et al*., 2007; Samaha *et al*., 2010), whereas the maturation of 25S involves a specific HID2 snoRNA (Zhu *et al*., 2016). The majority of plant snoRNAs are transcribed in polycistronic clusters (Rodor *et al*., 2010) and mature species are released by endo- and exoribonucleolytic processing. To check whether lack of DXO1 affects the levels of HID2, U3, U14 or other snoRNAs and in this indirect way impacts pre-rRNA maturation, we performed northern blot analysis for HID2, snoR10, U3 and U14 snoRNAs and, as controls, U2 and U4 snRNAs. As expected, the level of U3 snoRNA was not changed in *dxo1-2* mutants since U3 in plants is transcribed from an independent monocistronic unit by Pol III (Rodor *et al*., 2010). In turn, HID2, U14 and snoR10 levels were only slightly decreased (Supplementary Fig. S2E). These modest effects are unlikely to lead to pre-rRNA processing defects caused by the *dxo1-2* mutation, but together with the altered expression of RPs may to some extent contribute to the observed delay in rRNA synthesis.

Taken together, our results suggest that DXO1 has an indirect effect on pre-rRNA processing, which is most likely caused by its role in regulating the expression of ribosomal proteins and processing factors. We envisage that even modest changes in the amount of these essential proteins caused by the lack of DXO1 could additively produce the observed effects.

### DXO1 affects the maturation of chloroplast rRNAs

Although DXO1 has not been detected in chloroplasts, its contribution to the functioning of this organelle is indicated by the plant-specific features of Arabidopsis DXO1 protein, pale-green pigmentation of DXO1-deficient plants associated with reduced chlorophyll b content and finally RNA-seq analyses of *dxo1-2* plants (Kwasnik *et al*., 2019). Given the processing defects of nuclear-encoded pre-rRNAs and changes in the expression of genes encoding chloroplast RPs in *dxo1-2* mutants, it seemed possible that DXO1 could also affect rRNA biogenesis in chloroplasts. To verify this hypothesis we tested the accumulation of chloroplast rRNA precursors in the *dxo1-2* mutant line by northern blot analysis. Indeed, we detected elevated levels of specific pre-rRNAs in the mutant (Fig. 3). The most pronounced forms were the 1.9 kb precursor containing 16S rRNA detected by probes P_c_1 and P_c_2, and the 3.2 kb precursor containing 23S and 4.5S rRNAs detected by probes P_c_3, P_c_4 and P_c_5. Also, a slight increase of the primary 7.4 kb transcript, which is hardly visible in wild-type plants, was observed in the *dxo1-2* line. As with nuclear-encoded rRNAs, the levels of mature 16S and 23S rRNAs remained unaffected (Supplementary Fig. S3B). Nevertheless, defects in rRNA processing in *dxo1-2* plants may still affect ribosome assembly, as these processes occur simultaneously (Peña *et al*., 2017).

**Figure 3.**
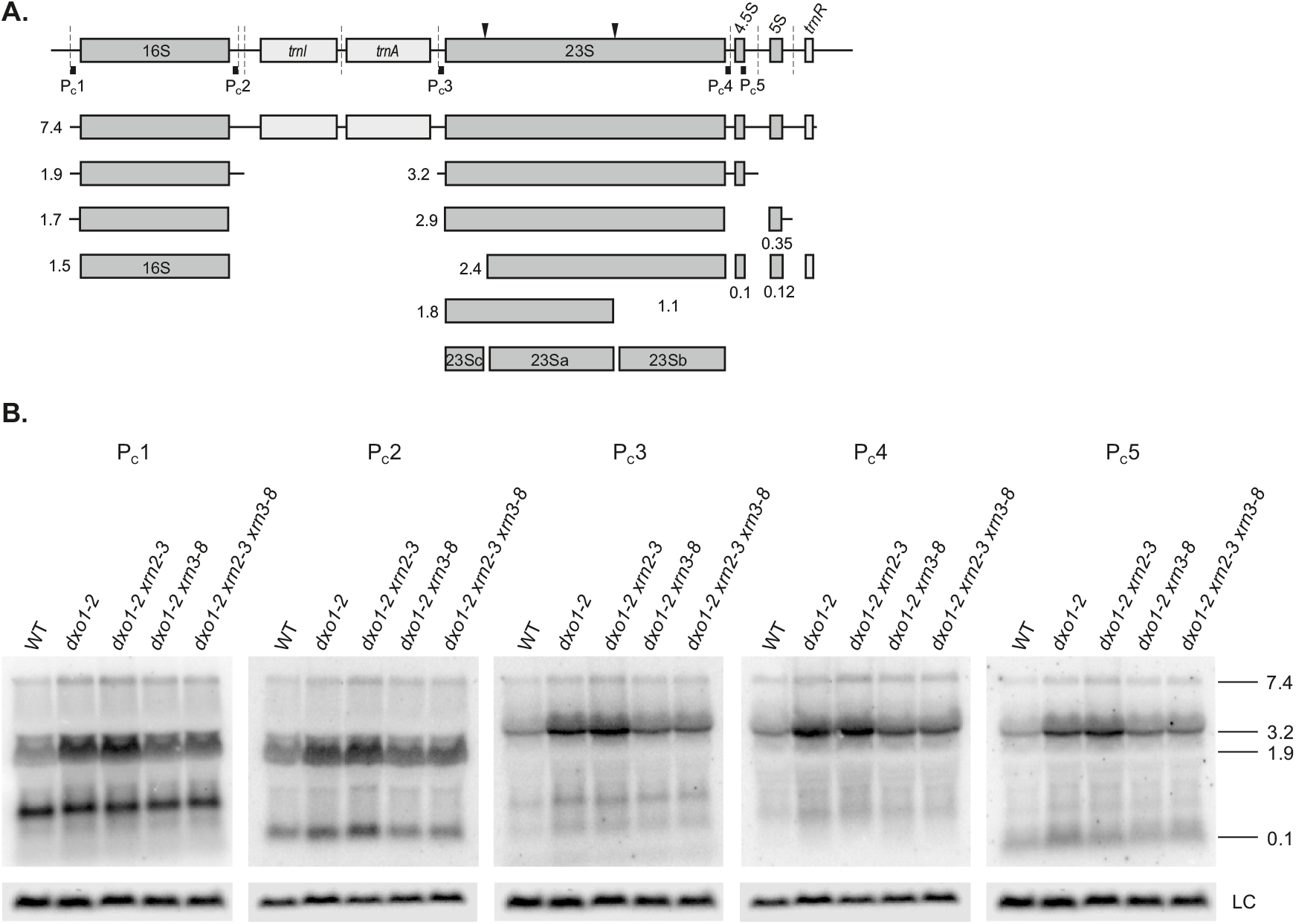
Chloroplast pre-rRNA processing defects caused by DXO1 knock-down are suppressed by *XRN3* silencing. (A) Diagram showing chloroplast rRNA precursor and processing intermediates. The location of probes used for Northern blot analysis is marked. (B) Northern blot analysis of chloroplast rRNA precursors and intermediates. Total RNA was extracted from 14-day-old WT and mutant seedlings shown, separated on 1.1% agarose gels and hybridized with probes specific for particular regions of pre-rRNA shown in (A). U2 snRNA was used as a loading control (LC). The length of specific precursors (kb) is marked on the right. Sequences of hybridization probes are listed in Supplementary Table S2.

Some accumulation of chloroplast pre-rRNAs was also visible in *dxo1-2 xrn2-3* and *dxo1-2 xrn2-3 xrn3-8* lines, but the *xrn3-8* mutation in the *dxo1-2* background reduced this effect (Fig. 3B). In contrast, there was no increase in chloroplast pre-rRNAs in single *xrn2-3* and *xrn3-8* or double *xrn2-1 xrn3-3* lines (Supplementary Fig. S3A). This observation, which supports the notion of the functional connection between DXO1 and XRN3 enzymes, is in contrast to the processing of nuclear-encoded pre-rRNAs, for which no phenotypic suppression was seen. Since neither of the proteins is present in chloroplasts, their possible interdependence is most likely related to the regulation of expression of nuclear-encoded factors involved in chloroplast rRNA biogenesis. According to RNA-seq analysis, nuclear-encoded chloroplast RPs are overrepresented in the group of genes that are upregulated in the *dxo1-2* mutant but downregulated in *xrn3-8* and *dxo1-2 xrn3-8* lines (Fig. 1H). These two opposing effects may be responsible for the complementation of the chloroplast rRNA maturation defects in the double *dxo1-2 xrn3-8* mutant. In turn, much less abundant chloroplast-encoded RP genes are mainly downregulated in *dxo1-2* and *dxo1-2 xrn3-8* lines (Fig. 1H). This profile is more common and, in addition to RP proteins, also a large part of other chloroplast-encoded mRNAs shows a decreased expression, most likely reflecting general chloroplast dysfunction (Supplementary Fig. S3C).

### Plant-specific N-terminal extension is required for DXO1 function in the processing of nuclear-encoded and chloroplast pre-rRNAs

The Arabidopsis DXO1 protein has several plant-specific features that influence its biochemical properties *in vitro* (Kwasnik *et al*., 2019). These include the asparagine residue (Asn298) located in the proximity of the DXO1 active site and a large and disordered N-terminal extension (NTE). Both elements, absent in other biochemically active DXO homologs, limit DXO1 enzymatic activity (Kwasnik *et al*., 2019).

To assess the contribution of the catalytic activity and plant-specific NTE to the function of DXO1 in pre-rRNA processing, we used transgenic *dxo1-2* lines expressing DXO1(WT), DXO1(E394A/D396A), DXO1(ΔN194) and DXO1(ΔN194/E394A/D396A) (Kwasnik *et al*., 2019). Expression of DXO1(WT) and catalytically inactive DXO1(E394A/D396A) variant, but not N-terminally truncated DXO1(ΔN194) and DXO1(ΔN194/E394A/D396A) forms, in the *dxo1-2* background restores the wild-type morphology and complements molecular phenotypes associated with changes in the levels of several mRNAs and RNA quality control siRNAs (rqc-siRNAs) (Kwasnik *et al*., 2019). Consequently, a similar pattern was observed for the processing of nuclear-encoded (Fig. 4A) and chloroplast pre-rRNAs (Fig. 4B). Northern blot analysis showed that the wild-type levels of rRNA precursors were restored in *dxo1-2* lines expressing the full-length DXO1, whereas accumulation of pre-rRNAs persisted in plants expressing the N-terminally truncated variants.

**Figure 4.**
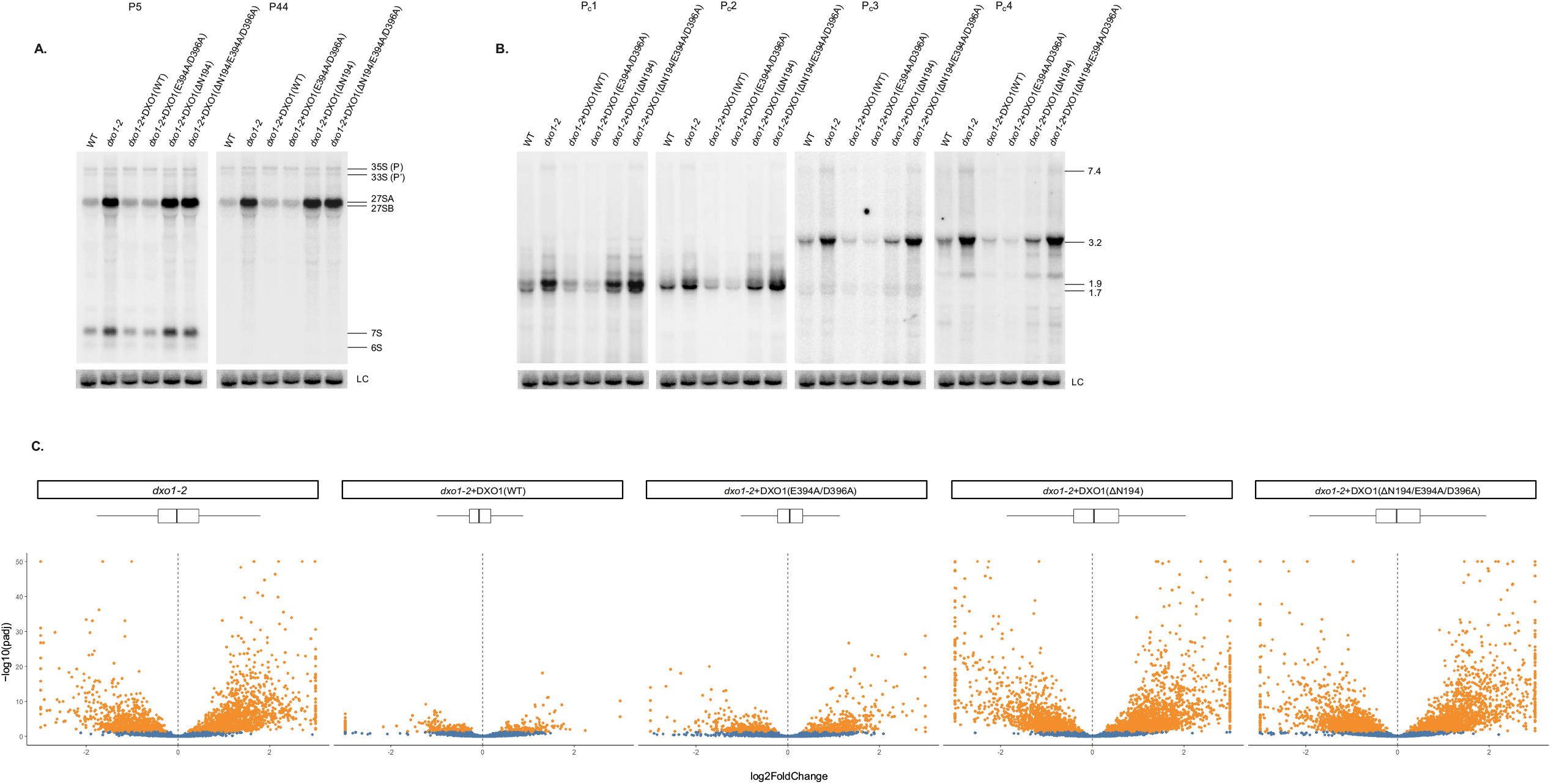
Molecular phenotypes of *dxo1-2* plants are rescued by the expression of the full-length WT or catalytically inactive DXO1, but not DXO1 variants lacking NTE. (A-B) Northern blot analysis of nucleolar (A) and chloroplast (B) pre-rRNAs in WT and *dxo1-2* mutant expressing DXO1(WT) or mutated variants: DXO1(E394A/D396A) catalytic mutant, DXO1(ΔN194) lacking the N-terminal domain and DXO1(ΔN194/E394A/D396A). eIF4A1 mRNA was used as a loading control (LC). Particular pre-rRNAs are described on the right. (C) Differential transcriptomic analysis of WT, *dxo1-2* and transgenic lines by 3’RNA-seq. Plots represent DESeq2 results with padj < 0.05 genes indicated in orange. Genes with differential expressions exceeding the x-axis limits were squished.

To confirm that the contribution of DXO1 to pre-rRNA processing is indirect and most likely mediated by changes in the expression of proteins involved in these pathways, we checked transcriptomic changes in all *dxo1-2* transgenic lines by 3’RNA-seq (Alpern *et al*., 2019). These data revealed that a large number of genes differentially expressed in *dxo1-2* was significantly decreased in lines expressing DXO1(WT) and DXO1(E394A/D396A), but not in those with DXO1 variants lacking NTE (Fig. 4C and Supplementary Table S3).

The analysis of intersections among these sets of genes showed the most significant overlap between *dxo1-2* and lines expressing DXO1(ΔN194) and DXO1(ΔN194/E394A/D396A) transgenes (Supplementary Fig. S4A). Consequently, gene clustering analysis showed that the expression level of most of the genes upregulated in *dxo1-2* was restored in *dxo1-2* transgenic lines transformed by full-length DXO1, either WT or with catalytic mutation, but not in lines with DXO1 ΔN variants (Supplementary Fig. S4B). These results show that, as with other phenotypes studied so far, the plant-specific N-terminal extension is also crucial for DXO1 function in pre-rRNA processing and this function does not depend on the enzymatic activity of DXO1. This raises the question of how important the enzymatic activity of DXO1 is for plant RNA metabolism and in what specific processes this activity plays a key role.

### DXO1 affects plant sensitivity to abiotic stress and temperature-related pre-rRNA processing

Increasing evidence indicates that the nucleolus, in addition to being a site for pre-rRNA processing and ribosome biogenesis, also plays an important role in adaptation to stress (Kalinina *et al*., 2018). Many different stressors affect the synthesis of rRNA and pre-rRNA processing factors and trigger changes in the pre-rRNA processing pathway (Palm *et al*., 2019; Sáez-Vásquez and Delseny, 2019; Martinez-Seidel *et al*., 2020). With regard to ribosome biogenesis, sugar is particularly important as it has been shown to promote rRNA biogenesis and to increase the expression of nucleolin, ribosomal proteins and components of the t-UTP pre-rRNA processing complex (Kojima *et al*., 2007; Ishida *et al*., 2016).

Among DEGs in the *dxo1-2* mutant, we found many genes involved in stress responses (Supplementary Table S1). To check whether defective pre-rRNA processing resulting from the *dxo1* mutation had an impact on the abiotic stress response, we performed growth tests of wild-type and *dxo1-2* plants in the presence of different concentrations of NaCl or glucose or in the absence of sugar. The effects of stress on pre-rRNA processing were assessed by northern analyses, also for drought, wounding and high temperature. The *dxo1-2* mutant showed a clearly increased sensitivity to salt stress and the lack of sugar, but grew better than wild-type seedlings in glucose-containing media (Fig. 5A). However, at the molecular level, we did not observe striking differences in pre-rRNA processing or accumulation of specific precursors in the mutant and/or wild-type in all stress conditions tested (Fig 5B and C). Induction of stress response to wounding, drought and high temperature was confirmed by checking the levels of known stress markers (Supplementary Fig. S5).

**Figure 5.**
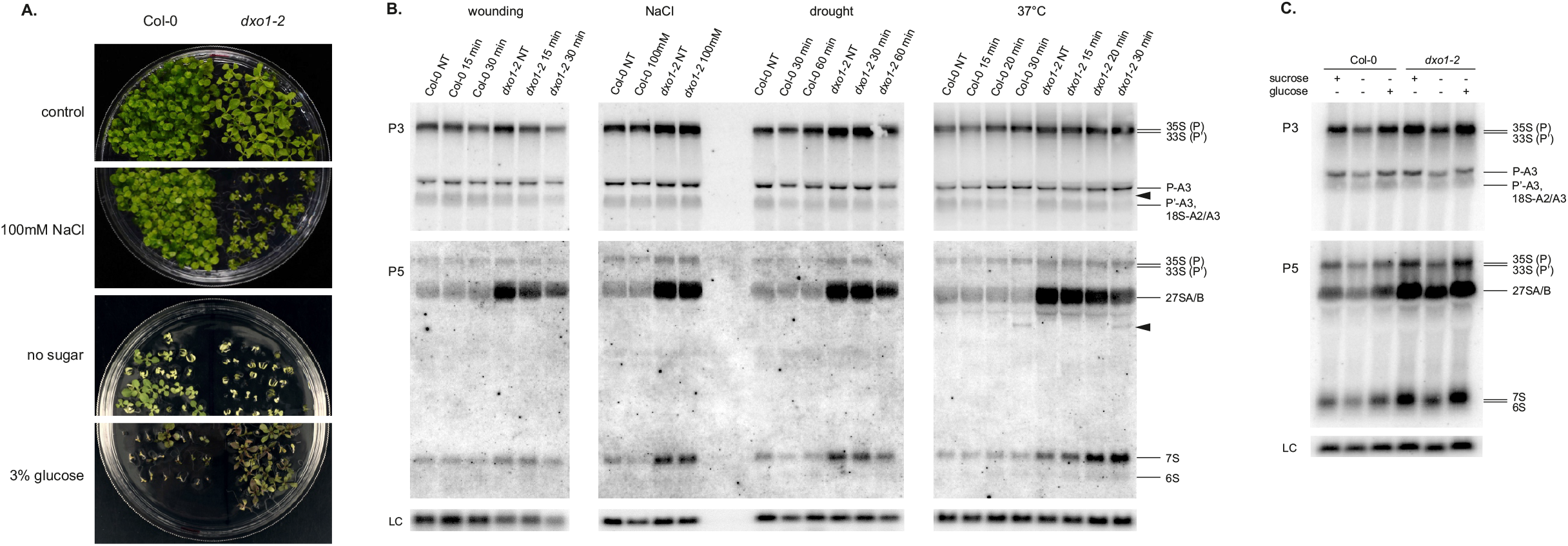
*dxo1-2* mutant seedlings respond to stress differently than WT. (A) 14-day-old seedlings of the WT and *dxo1-2* mutant grown under regular growth conditions (MS medium containing 1% (w/v) sucrose; control), on MS medium without any sugar added (no sugar), on MS medium with 100 mM NaCl and 1% (w/v) sucrose or on MS medium containing 3% glucose (instead of sucrose). (B and C) Northern blot analysis of pre-rRNAs in WT and *dxo1-2* seedlings treated by wounding, drought or high temperature (37°C) for the indicated time (minutes) or grown on plates containing 100 mM NaCl for high salinity stress (B) or grown on plates with different sugar conditions (C). Total RNA was extracted from 14-day-old seedlings treated with the indicated stressors, resolved on 1.1% agarose gels and hybridized with probes P3 and P5. Precursors and rRNA processing intermediates are described on the right. Black arrowheads indicate the P-C2 intermediate, symptomatic of the ‘ITS2-first’ processing pathway. U2 snRNA was used as a loading control (LC).

Interestingly, while we did not detect *dxo1-2* mutant-specific effects of stress on pre-rRNA processing, we did observe some changes in both the wild-type and the mutant subjected to heat treatment (37°C) for up to 30 minutes. In particular, heat-stressed plants showed a gradual decrease of the P’-A3 precursor (Fig. 5B; probe P3) and 27S variants (Fig. 5B; probe P5). We also detected an additional precursor visible after 30 minutes at 37°C (Fig. 5B; probe P5, marked with an arrowhead). This species most likely corresponds to a stress-related P-C2 intermediate that was initially identified in *irp7* mutants upon auxin treatment and represents a bypass pre-rRNA processing pathway called ‘ITS2-first’, in which the initial cleavage of the primary precursor occurs at site C2 within ITS2 (Palm *et al*., 2019; Sáez-Vásquez and Delseny, 2019; Shanmugam *et al*., 2021; Darriere *et al*., 2022).

These preliminary analyses clearly show that although DXO1 is required for abiotic stress response, as was previously reported in the case of abscisic-acid (Yu *et al*., 2021), this functional dependency is not related to its contribution to pre-rRNA biogenesis.

## DISCUSSION

### The contribution of DXO1 to pre-rRNA processing

The role of DXO1 in ribosome biogenesis is manifested through the morphological phenotype of *dxo1-2* plants that resemble other known *Arabidopsis* ribosome-related mutants, strong upregulation of RP genes and accumulation of rRNA precursors. This function of DXO1 would not be surprising, as fungal Rai1 and Dxo1 proteins have already been shown to directly participate in pre-rRNA processing, albeit in the case of Rai1 this activity requires interaction with other proteins (Rat1, Las1-Grc3) (Fang et al., 2005; Gasse et al., 2015; Hurtig and van Hoof, 2022; Xue et al., 2000). Ribosome biogenesis in plants is much less understood than in other kingdoms, and only a few plant enzymes and factors involved in pre-rRNA processing have been identified. We show that DXO1 not only contributes to the biogenesis of cytosolic ribosomes but also has an impact on the maturation of chloroplast rRNAs. However, our data strongly suggest that the role of DXO1 in these processes is most likely indirect. First of all, the nuclear and chloroplast pre-rRNA processing defects observed in *dxo1* plants are independent of DXO1 enzymatic activity. Instead, the correct pre-rRNA maturation pathway requires the presence of the plant-specific N-terminal extension. Moreover, although DXO1 affects pre-rRNA processing in the nucleus and chloroplasts, it is localized to the nucleus and, to a lesser extent, the cytoplasm, but not to chloroplasts (Kwasnik *et al*., 2019). On the other hand, the lack of DXO1 causes changes in the expression of both cytoplasmic and organellar RP genes. Taken together, these observations suggest that pre-rRNA processing defects in the *dxo1-2* mutant are a secondary effect resulting from the deregulation of RP protein expression, which consequently leads to deficient ribosome biogenesis and assembly. The contribution of RPs to pre-rRNA processing, pre-ribosome assembly and export has been demonstrated in many cases in yeast and mammalian cells (Sloan *et al*., 2016). Alternatively, DXO1 may recruit other pre-rRNA processing factors, possibly via its plant-specific unstructured N-terminal domain, which is required for DXO1 cellular functions (Kwasnik et al., 2019 and this work).

Unexpectedly, *dxo1-2* mutants accumulate 3’-extended, but not 5’-extended, precursors of 5.8S rRNA. This observation supports the notion that DXO1 does not act as a 5’-3’ exoribonuclease and that its residual activity observed *in vitro* is probably not relevant for its function *in vivo*. This is in contrast to effects reported in yeast for mutants in 5’-3’ exoribonucleases, Rat1 and Rrp17, or Rai1, where maturation of both 5’- and 3’-ends of 5.8S was impaired (Fang et al., 2005; Oeffinger et al., 2009). It has even been suggested that yeast Rai1 may coordinate 5’ and 3’ processing by Rat1 and Rrp6, respectively. It turns out that plant DXO1 is distinct from homologs in other organisms, mainly due to plant-specific features that alter its biochemical properties and, consequently, cellular functions (Kwasnik *et al*., 2019). There are also other DXO family proteins with a very low enzymatic activity, e.g. *Candida albicans* Rai1 (Wang *et al*., 2015), or even catalytically inactive, like *Drosophila melanogaster* Cutoff, which is involved in piRNA synthesis by preventing premature transcription termination (Mohn *et al*., 2014; Chen *et al*., 2016; Andersen *et al*., 2017). Thus, the wide repertoire of functions represented by DXO/Rai1 proteins varies greatly between organisms and is not necessarily associated with enzymatic activities.

The plant-specific N-terminal extension of the DXO1 protein may also play a role in the formation of the nucleolar structure, which in turn may influence pre-rRNA processing and ribosome assembly. The nucleolus is one of the membrane-free biomolecular condensates (liquid droplets), which is sequestered from the nucleoplasm through liquid-liquid phase separation (LLPS) (Yoneda *et al*., 2021). LLPS of the nucleolus depends on the interactions of liquid droplet-forming components, including DNA, RNA and proteins. The concentration of factors, enzymes and nucleic acids in a condensate facilitates nucleolar functions such as pre-rRNA processing and the formation of ribosomal subunits (Yoneda *et al*., 2021). One of the driving forces behind LLPS formation are interactions mediated by intrinsically disordered protein regions (IDRs) (Emenecker *et al*., 2021). Since the N-terminal extension of DXO1 is intrinsically disordered (Kwasnik *et al*., 2019), DXO1 may impact ribogenesis by driving the formation of intra-nucleus droplets by LLPS, but more research is needed to investigate this possibility.

### The effect of stress on ribosome biogenesis

The functioning of both the nucleolus and chloroplasts is closely related to the plant’s response to stress. Chloroplasts act as sensors to receive and transmit stress signals to the cell nucleus via the so-called retrograde signaling pathway (Mielecki *et al*., 2020). In turn, several stresses affect the nucleolar structure and protein composition (Kalinina *et al*., 2018). It is believed that the existence of alternative pre-rRNA processing pathways and redundant enzymes in the nucleolus secure the production of functional ribosomes in adverse conditions. Accordingly, several Arabidopsis pre-rRNA processing mutants exhibit altered sensitivity to various stressors (e.g. *irp* (Palm *et al*., 2019), *apum23* (Huang *et al*., 2018), *apum24* (Maekawa *et al*., 2018)) and the plant response to abiotic stresses includes altered expression of pre-rRNA processing factors and changes in the pre-rRNA maturation pathway (Palm *et al*., 2019; Sáez-Vásquez and Delseny, 2019; Martinez-Seidel *et al*., 2020; Shanmugam *et al*., 2021; Darriere *et al*., 2022). In response to stress, the expression of RP genes is regulated, inter alia, by the TOR signaling pathway (Petibon *et al*., 2020). This pathway may be impaired in the *dxo1-2* mutant because among genes downregulated in our RNA-seq data there are components of the TORC1 complex: TOR (Target of Rapamycin protein kinase, AT1G50030), RAPTOR (Regulatory-associated protein of TOR, AT3G08850) and LST8 (Lethal with Sec Thirteen 8, AT3G18140), and several factors of the TOR signaling pathway, including S6K2 (ribosomal protein S6 kinase 2, AT3G08720) (Supplementary Fig. S6). The TOR kinase is a highly conserved key regulator of growth and development that responds to nutritional and stress signals and controls ribosome abundance (Wullschleger *et al*., 2006; Deprost *et al*., 2007; Ren *et al*., 2012; Wang *et al*., 2018; Petibon *et al*., 2020; Scarpin *et al*., 2020; Martínez-Pacheco *et al*., 2021; Busche *et al*., 2021). Importantly, the *Arabidopsis* TOR protein has been shown to control pre-rRNA synthesis through direct interaction with the promoter and 5’ETS in rDNA (Ren *et al*., 2011) and to regulate transcription and translation of RPs (Dobrenel *et al*., 2016; Scarpin *et al*., 2020). It is tempting to speculate that DXO1 regulates the levels of the TORC1 components, and as a result the lack of DXO1 affects ribosome biogenesis. This is further supported by the altered response of *dxo1-2* plants to nutrient availability (high sugar and sucrose starvation). Still, besides TOR signaling, there are other mechanisms by which plant stress responses regulate ribosome formation and function (Sáez-Vásquez and Delseny, 2019; Martinez-Seidel *et al*., 2020).

Environmental stress, especially nutrient availability and heat, has been shown to alter pre-rRNA processing in Arabidopsis (Maekawa *et al*., 2018; Shanmugam *et al*., 2021; Darriere *et al*., 2022). Indeed, in plants treated with elevated temperatures we observed changes in the accumulation of some rRNA precursors and detected an unusual P-C2 intermediate encompassing the 5.8S and 18S rRNAs. These observations are consistent with recent studies on rRNA maturation under heat stress in plants, reporting an altered abundance of individual pre-rRNAs at low and/or high temperatures (Hang *et al*., 2018; Liu *et al*., 2020; Shanmugam *et al*., 2021; Darriere *et al*., 2022). These changes may reflect the influence of environmental temperature on plant ribosome biogenesis in natural habitats. The P-C2 species resembles the atypical intermediates detected in yeast (24S) and human cells (30SL3’ and 21SL3’) lacking the RRP5 processing factor, which result from blocking the initial processing steps within 5’ETS and ITS1 spacers (Venema and Tollervey, 1996; Sloan *et al*., 2013). It is possible that in heat-stressed plants pre-rRNA maturation, particularly at the initial stages, is slowed down or arrested, and this gives rise to the P-C2 intermediate. Alternatively, elevated temperature can trigger the atypical ‘ITS2-first’ pre-rRNA processing pathway (Palm *et al*., 2019; Sáez-Vásquez and Delseny, 2019). Nevertheless, since the absence of DXO1 has no effect on pre-rRNA processing in stressed plants, the altered stress sensitivity of the *dxo1-2* mutant is not related to rRNA maturation. In turn, changes in the level of RP mRNAs, followed by their modified translation under stress, may affect the mutant response to stress due to the adaptation of the ribosome composition to environmental conditions (Kawaguchi *et al*., 2004; Nicolaï *et al*., 2006; Hummel *et al*., 2012; Martinez-Seidel *et al*., 2020).

### Functional interconnection between DXO1 and XRN proteins

The partial suppression of *dxo1-2* phenotypes by mutations in *XRN2* and/or *XRN3* genes and those of *xrn3-8* by the lack of DXO1 points to the possible functional interaction between DXO1 and XRN proteins, XRN3 in particular. However, these connections appear very complex and concern specific aspects of the cellular activities of both proteins. The *xrn2-3* mutation reverts only the growth retardation and pointed-leaf phenotype of *dxo1-2* plants, while *XRN3* silencing also restores normal processing of chloroplast pre-rRNAs. In addition, *XRN*-mediated suppression does not concern the pale green coloration of the mutant or defects in nuclear pre-rRNA processing. These observations indicate that the contribution of DXO1 to the maturation of cytosolic or chloroplast rRNAs has little impact on severe morphological or molecular aberrations caused by the *dxo1* mutation. Therefore, partial inversion of some of the *dxo1* phenotypes by *XRN3* knock-down is most likely due to attenuation of transcriptomic changes caused by DXO1 deficiency.

The most interesting case is represented by the chloroplast-related processes. The fact that the pale-green pigmentation phenotype of *dxo1* plants is not rescued by *xrn* mutations, especially *xrn3-8*, is consistent with a similarly reduced expression of chloroplast genes in single *dxo1-2* and double *dxo1-2 xrn3-8* lines. In fact, the downregulation of these genes is even more pronounced in *dxo1-2* with silenced *XRN3* (see figures 1H and S3C), suggesting an additive effect. Such profiles reflect general chloroplast dysfunction, which is arguably even stronger in plants lacking both proteins. The biochemical activity of XRNs is connected with chloroplast function via the 3′-phosphoadenosine 5′-phosphate (PAP) signaling pathway (Estavillo *et al*., 2011), as is likely the case of DXO1, which is also inhibited by PAP (Kwasnik *et al*., 2019). PAP regulates nuclear gene expression in response to high light and drought stress, possibly by inhibiting 5′-3′ RNA degradation by XRNs (Estavillo *et al*., 2011). It is therefore feasible that both DXO1 and XRNs are controlled by a mechanism mediated by PAP regulators such as FRY1, a nuclear-encoded protein located in chloroplasts and mitochondria that dephosphorylates PAP (Estavillo *et al*., 2011). Interestingly, FRY1 has also been shown to repress the biogenesis of ribosomal RNA-derived siRNAs (risiRNAs) from the rDNA loci that arise from aberrant rRNA precursors, mainly due to the dysfunction of 5’-3’ exoribonucleases (You *et al*., 2019). These observations point to the existence of an autoregulatory feedback loop in which DXO1 and XRNs regulate chloroplast retrograde signaling, which in turn affects their activity.

## SUPPLEMENTARY DATA

**Figure S1**. (A) Phenotypes of mutant lines presented in this study. (B) Gene clustering analysis of RNA-seq data.

**Figure S2**. Northern blot analysis of pre-rRNAs (A), mature rRNAs (C and D) and snoRNAs (E) in *dxo1-2, xrn* mutants and double and triple mutant lines.

**Figure S3**. (A-B) Northern blot analysis of chloroplast pre-rRNAs and rRNAs.(C) Scatter plot of RNA-seq data for genes encoded in chloroplasts.

**Figure S4**. 3’RNA-seq analysis of WT, *dxo1-2* and *dxo1-2* transgenic lines expressing DXO1(WT), DXO1(E394A/D396A), DXO1(ΔN194) or DXO1(ΔN194/E394A/D396A).

**Figure S5**. Northern blot analysis of known stress marker mRNAs in WT plants and the *dxo1-2* mutant.

**Figure S6**. A diagram and table showing transcriptomic changes in the *dxo1-2* mutant of factors involved in the TOR signaling network, according to RNA-seq data.

**Table S1**. List of genes included in clusters presented in Figure 1G and Supplementary Figure S1B-C.

**Table S2**. Oligonucleotides used in this study.

**Table S3**. Number of differentially expressed genes in 3’RNA-seq analysis and list of genes included in clusters presented in Supplementary Figure S4B.

## AUTHOR CONTRIBUTIONS

MZ-P developed the research concept and wrote the manuscript with the contribution of all authors; MZ-P, AK, MK and AG-M performed the experiments; AK and MK analyzed the RNA-seq and 3’RNA-seq data; JK supervised and completed the writing; all authors read and approved the manuscript.

## DATA AVAILABILITY

The data that support the findings of this study are openly available; RNA-seq data are deposited in the Gene Expression Omnibus database under accession codes GSE95473 (Col-0 and *xrn3-8* plants), GSE99600 (*dxo1-2* plants) and GSE210614 (*dxo1-2 xrn3-8* double mutant). 3’RNA-seq data are deposited upon GSE210631.

## FIGURE LEGENDS

**Supplementary Figure S1.** (A) The double mutant lines *dxo1-2 xrn2-3* and *dxo1-2 xrn3-8* and the triple mutant *dxo1-2 xrn2-3 xrn3-8* present less severe phenotypes than the single *dxo1-2* mutant. Plants were grown in soil in Percival growth chambers at 22°C on a 16-h-light/8-h-dark photoperiod and pictures were taken after 21 days (21D) and 28 days (28D) of growth. (B-C) Gene clustering analysis of RNA-seq data from *dxo1-2, xrn3-8* and *dxo1-2 xrn3-8* double mutant, for genes upregulated (B) and downregulated (C) in the *dxo1-2* line. Clustering was performed using standard R functions. Genes included in each cluster and GO term enrichment analysis of each cluster are listed in Supplementary Table S1. The GO term Overrepresentation Test revealed the enrichment of GO categories related to the following processes: (B) cluster 3, maturation of LSU-rRNA, translation and response to light; cluster 4, translation, ribosomal SSU assembly and LSU biogenesis; rRNA processing, chlorophyll metabolic processes and response to light; cluster 5, regulation of transcription and RNA processing; cluster 6, RNA methylation; rRNA processing, protein complex assembly, translation, transport, carbohydrate metabolic process; developmental growth, defense response; clusters 7-9, cell division and development; cluster 10, response to stimulus, cluster 11, cell division; cluster 13, ncRNA processing and RNA modification, embryo development; cluster 15, circadian rhythm, photoperiodism, flowering, stress responses, development; (C) clusters 1-4, stress and defense responses, cell communication, signaling pathways and metabolic processes; cluster 5, cytosolic transport, signal transduction and protein localization; cluster 6, stress and defense responses, photosynthesis; cluster 7, defense responses, signal transduction and localization; cluster 8, development, intracellular transport, protein localization; cluster 9, regulation of transmembrane transport; cluster 10, stress and defense responses; clusters 11-12, response to stimulus; cluster 13, plant organ senescence, defense response; clusters 15-16, stress and defense responses and metabolic processes.

**Supplementary figure S2.** Northern blot analysis of pre-rRNAs (A), mature rRNAs (C and D) and snoRNAs (E) in *dxo1-2* and *xrn* mutants and double and triple mutant lines. Total RNA was extracted from 14-day-old seedlings. RNA was separated on 1.1% agarose (A and C) or 6% polyacrylamide (D and E) gels and hybridized with probes specific for particular regions of pre-rRNA depicted in (A) or with probes specific to the indicated rRNAs and snoRNAs (C-E). Hybridizations for U2 snoRNA were used as loading controls (LC). On (A) pre-rRNAs and rRNA processing intermediates are described on the right. White arrowheads indicate the 5’ETS-A3 intermediate. Black arrowheads correspond to 27SA pre-rRNA. White asterisks indicate P-A3 and black asterisks denote P’-A3 and 18S-A2/A3 precursors. Arrows indicate by-products that accumulate in *xrn* mutants: black arrows show fragments of 5’ETS ending at the site P’ (5’ETS-P’) and white arrow marks the 5’ETS-P1 by-product. (B) Primer extension analysis of 25S 5’ end in wild-type (WT) and *dxo1-2* mutants, performed using primer 25S-5’. Sequences of hybridization probes are listed in Supplementary Table S2.

**Supplementary figure S3.** Northern blot analysis of chloroplast pre-rRNAs (A) and mature chloroplast rRNAs (B) in *dxo1-2* and indicated mutant lines. Total RNA was extracted from 14-day-old seedlings. RNA was separated on 1.1% agarose gels and hybridized with probes specific for particular regions of pre-rRNA depicted in (A) or with probes specific for the indicated mature rRNAs (B). U2 snRNA was used as a loading control (LC). The length of specific precursors (kb) is marked on the right. Sequences of hybridization probes are listed in Supplementary Table S2. (C) Scatter plot of genes encoded in chloroplasts. Orange dots represent differentially expressed genes (padj < 0.05) and blue dots represent non-differentially expressed genes. 3’RNA-seq analysis of WT, *dxo1-2* and *dxo1-2* transgenic lines expressing DXO1(WT), DXO1(E394A/D396A), DXO1(ΔN194) or DXO1(ΔN194/E394A/D396A). (A) Overlaps between the upregulated and downregulated genes in the indicated data sets, represented with UpSet plots that were generated using the UpSetR package. (B) Gene clustering analysis of 3’RNA-seq results performed for all DEGs in indicated plant lines. Hierarchical clustering on mean scaled read counts was used to obtain differential expression patterns across samples. Parallel coordinate plots were generated with ggparcoord and grouped by cluster.

Northern blot analysis of known stress marker mRNAs in WT plants and the *dxo1-2* mutant. Total RNA was extracted from 14-day-old seedlings treated with the selected stressors for the indicated time (minutes), resolved in 1.1% agarose gels and hybridized with probes specific for denoted mRNAs. U2 snRNA was used as a loading control (LC).

**DXO1 dysfunction deregulates the TOR signaling pathway.** A diagram and table showing transcriptomic changes in the *dxo1-2* mutant of factors involved in the TOR signaling network, according to RNA-seq data (Kwasnik *et al*., 2019). Genes with upregulated or downregulated expression in the *dxo1-2* mutant are marked in red or blue, respectively. Asterisks indicate genes with expression restored to the WT level in the double *dxo1-2 xrn3-8* mutant. The activity of the TORC1 complex is regulated by environmental cues, such as light, stress or nutrient starvation (e.g. carbon, amino acids or nucleotides). Assembly of the TORC1 complex, composed of the TOR, RAPTOR and LST8 proteins, which are downregulated in the *dxo1-2* mutant, is aided by assembly factors, many of which are upregulated in the *dxo1-2* (shown in the lower left corner). PRS4, cytosolic phosphoribosyl pyrophosphate (PRPP) synthetase; PRPP, a key metabolite in nucleotide biosynthesis, necessary for TOR activity and rRNA synthesis (Busche *et al*., 2021); S6Ks, p70 ribosomal S6 kinases, activated by binding of TORC1, known to phosphorylate RPS6A and RPS6B ribosomal proteins in a day- and night-dependent manner, as well as many translation factors (Obomighie *et al*., 2021). Translation factors differentially expressed in *dxo1-2* are shown on the rightside of the diagram. Arabidopsis TOR kinase domain directly regulates pre-rRNA synthesis by binding to the *45S rRNA* promoter and 5’ETS (Ren *et al*., 2011). The table on the right shows changes in expression (FC – fold change) of depicted genes in the *dxo1-2* mutant.

